# The Functional Role of Pinwheel Topology in the Primary Visual Cortex of High-Order Animals for Complex Natural Image Representation

**DOI:** 10.1101/2024.03.07.583885

**Authors:** Haoyu Wang, Haixin Zhong, Wei P Dai, Yuguo Yu

**Author notes:** Joint first authors (Haoyu Wang, Haixin Zhong).

## Abstract

The primary visual cortex (V1) of high-level animals exhibits a complex organization of neuronal orientation preferences, characterized by pinwheel structure topology, yet the functional role of those complex patterns in natural image representation remains largely unexplored. Our study first establishes a new self-evolving spiking neural network (SESNN) model, designed to mimic the functional topological structure of orientation selectivity within V1. We observe the emergence of a particularly new “spread-out” firing patterns from center to the surround of the pinwheel structures in response to natural visual stimuli in pinwheel structures, propagating from pinwheel centers and spreading to iso-orientation domains—a pattern not found in salt-and-pepper organizations. To investigate this phenomenon, we propose a novel deep recurrent U-Net architecture to reconstruct images from V1’s spiking activity across time steps and assess the encoded information entropy of different firing patterns via the model’s predicted uncertainty, offering a spatiotemporal analysis of V1’s functional structures. Our findings reveal a trade-off between visual acuity and coding time: the “spread-out” pattern enhances the representation of complex visual details at the cost of increased response latency, while salt-and-pepper organizations, lacking such domains, prioritize rapid processing at the expense of reduced visual acuity. Additionally, we demonstrate that this trade-off is modulated by the size of iso-orientation domains, with larger domains—supported by denser neuronal populations—substantially improving both visual acuity, coding efficiency, and robustness, features diminished in smaller domains and salt-and-pepper arrangements. Our research provides a foundational understanding of the principles underlying efficient visual information representation and suggests novel strategies for advancing the robustness and performance of image recognition algorithms in artificial intelligence.

## 1 Introduction

Research demonstrates that neurons in the primary visual cortex (V1) acquire orientation selectivity[3, 10, 11, 23, 29]. Notably, in species like primates, V1 is characterized by pinwheel structures, evident in systematic shifts in orientation preference maps (OPMs). Conversely, species such as rodents display more scattered salt-and-pepper organizations[9, 12, 14–16]. Yet, understanding of the functional dynamics within these structures, especially neuron responses in iso-orientation domains (IODs) and pinwheel centers (PCs), remains limited. This knowledge gap is crucial in understanding how neurons process external visual stimuli through these distinct structures across species. Addressing it is crucial to advance our knowledge of visual processing in the brain and to inform the creation of algorithms replicating these complex neural structures.

To investigate the functional significance of pinwheel structures in V1, we propose a groundbreaking hybrid model comprising a self-evolving spiking neural network (SESNN, detailed in section 2.2) and a deep recurrent U-Net (SESNN-DRUnet). (1) The SESNN module transforms natural images into spike patterns, simulating the development of OPMs through natural images. This approach, leveraging synaptic plasticity and natural images (section 2.1), enables the modeling of realistic pinwheel and salt-and-pepper patterns, capturing complex spatial-temporal responses within pinwheels - a feat not entirely feasible through experimental methods[25]. (2) The DRUnet module, detailed in section 2.3, decodes visual stimuli from V1 neuron spike data, focusing on firing rates and precise timing. This novel architecture is the first to capture pinwheels’ temporal-spatial structure at single time steps. We train the DRUNet component on our simulated V1 spike data to optimize visual reconstruction and quantify image uncertainty, reflecting the information entropy. The decoding accuracy is evaluated using metrics discussed in section 2.3.

The main contribution of our work can be summarized as three-fold:

- Research [25] shows that stimulating small areas within pinwheel structures triggers delayed responses elsewhere, with full neuronal recording within pinwheels being challenging. Our SESNN module addresses this by enabling the capture response of all neurons within OPMs. Moreover, the DRUnet module provides novel insights into the visual representation within pinwheel structures, exploring the intricate interplay of temporal and spatial coding.
- We demonstrate that firing waves originate in pin-wheel centers (PWCs), where diverse orientation preferences are represented, and then propagate to IODs for enhanced orientation-specific processing, facilitated by short-distance lateral connections between neurons within IODs.
- Our findings reveal a trade-off between response speed and visual acuity in pinwheel structures versus salt-and-pepper organizations. Pinwheel structures, indicative of advanced edge detection capabilities, feature densely packed neurons in PWCs that intricately capture visual stimuli. Although this results in enhanced orientation-specific details, it lengthens coding time. Conversely, the less neuron density and uncoordinated firing in salt-and-pepper organizations enable faster stimulus encoding but compromise visual acuity, leading to less precise visual information processing.

## 2 Methods

### 2.1 Stimulus images

In order to maximize the diversity of oriented visual inputs during training of the SESNN component, as well as enhance the model’s generalization ability for decoding,

We utilized a dataset of 20 original whitened images (512×512 pixels), each rotated clockwise in 5-degree increments to capture diverse orientation details. This rotation generated 72 variations per original image, amounting to a total of 1440 images in our training dataset. The images are resized to either 130×130 or 127×127 pixels via bilinear interpolation to accommodate receptive field (RF) sizes under different parameter arrangements. All images undergo equivalent whitening to eliminate pairwise correlations among pixels to emulate processing in the lateral geniculate nucleus (LGN) [30]. No additional enhancements or modifications are made to the raw image dataset.

During the self-organization of SESNN, each epoch consisted of excitatory neurons receiving a 100ms exposure to 1440 images randomly selected from the dataset, while inhibitory neurons received no visual stimulation. The RFs of different excitatory neurons exhibited overlapping coverage of the visual fields, with the degree of overlap varying to reflect species-dependent iso-orientation patch sizes in the visual cortex for a given retinal location (see section 2.2 for details).

For the training of the decoding model, a custom dataset was curated, comprising images from three distinct categories. Each category included 4,000 images: portraits from the CelebA dataset[17], automobile images from Stanford car dataset[5] and aircraft from Kaggle (Detailed information about the dataset is provided in the Supplementary Materials.). These images were presented to the SESNN module, generating spike trains with a duration of 100 milliseconds. The dataset was partitioned, allocating 80% for the training of the decoder, while the remaining 20% was reserved for the evaluation of decoding quality.

### 2.2 The SESNN module

The SESNN module (refer to Figure 2 a-e) is specifically designed to facilitate the self-organization of visual representations, which is instrumental in elucidating the functional roles of various orientation preference structures within the primary visual cortex across different species. It incorporates a network of excitatory neurons (E-neurons) and inhibitory neurons (I-neurons), maintaining a ratio of 4:1 (non-human primates) and 9:1 (rodents) between excitatory and inhibitory neurons [2]. All the neurons are spatially positioned on a two-dimensional lattice, with initial connection weights modeled by a Gaussian function of their distance,

**Figure 1.**
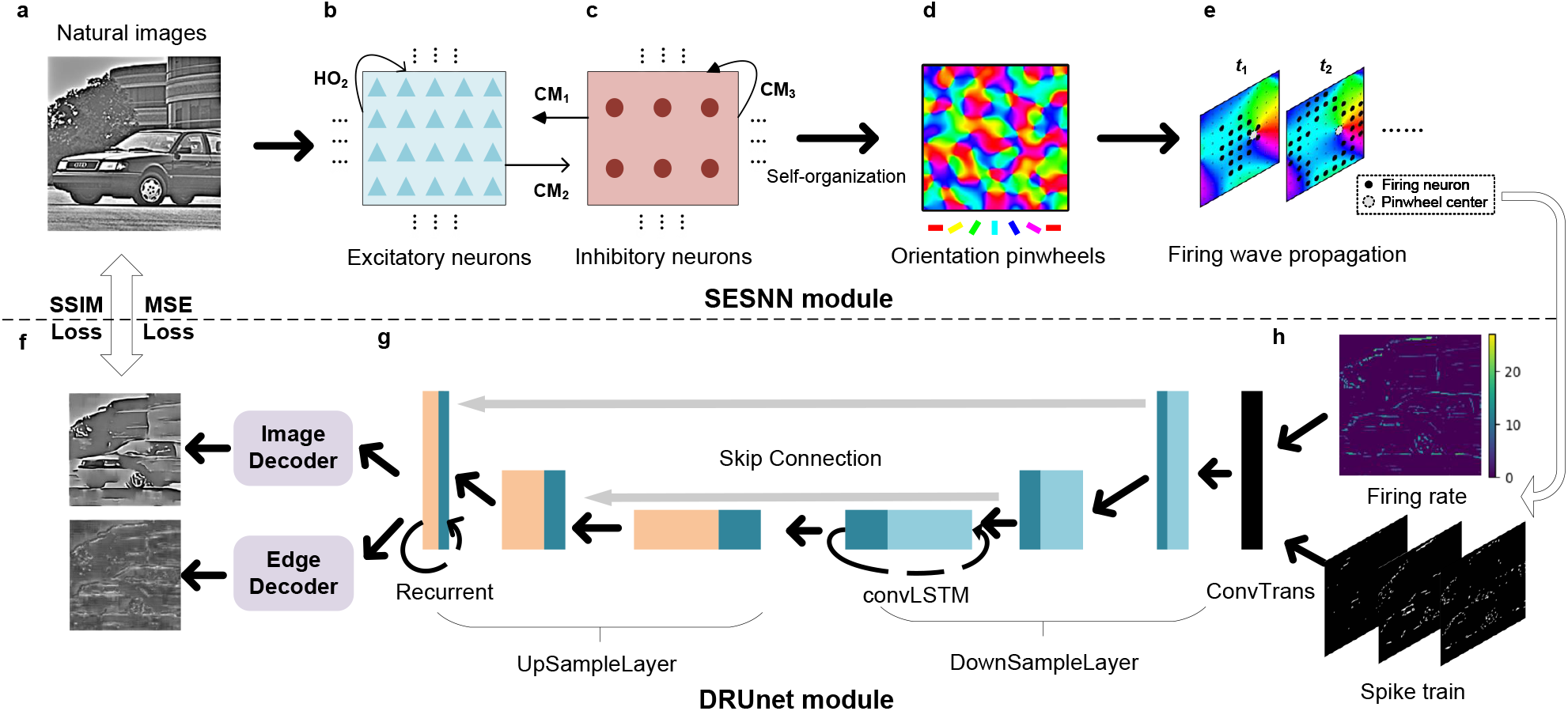
The framework of SESNN-DRUnet model. **a** The input to the SESNN module consists of natural images. **b** Excitatory neurons of SESNN. **c** Inhibitory neurons of SESNN. **d** The interactions between excitatory and inhibitory neurons lead to the self-organization of neurons into orientation pinwheels. **e** SESNN emerges orientation pinwheel structures. **f** This part employs Mean Squared Error (MSE) and Structural Similarity Index (SSIM) as loss functions to evaluate the reconstruction quality of the output against the input images, balancing fidelity and perceptual similarity. **g** This part is a deep recurrent neural network with convolutional LSTM layers and skip connections, suggesting a focus on capturing temporal dynamics and spatial hierarchies in the data for reconstruction purposes. **h** The outputs of the DRUnet module include a visualization of the firing rate and the extracted spike train data, which are indicative of the neural response patterns simulated by the model.

**Figure 2.**
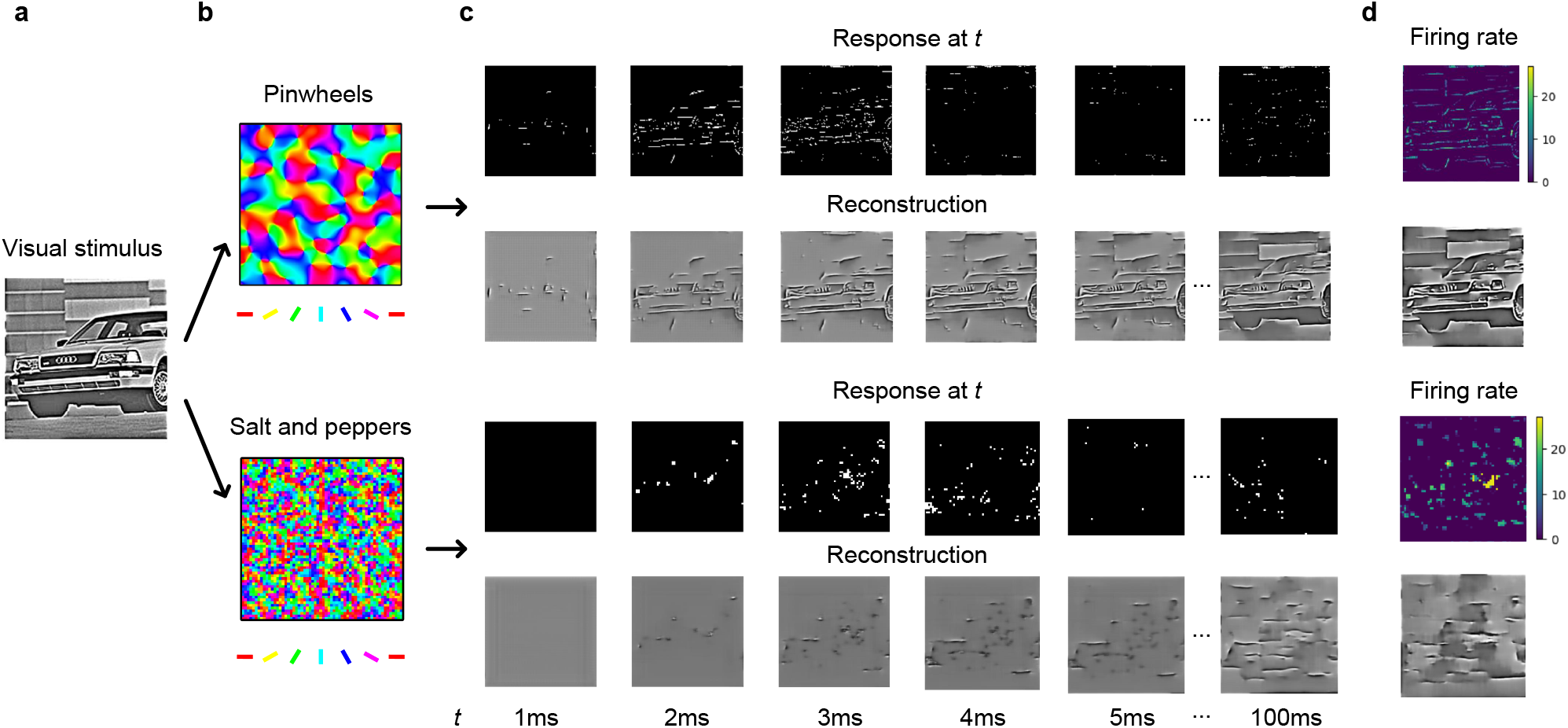
The comparative analysis of neural responses and image reconstruction quality between two types within V1: pinwheels and salt-and-pepper organizations. **a** The grayscale image of a car serves as the initial visual stimulus. **b** Two firing patterns of pinwheels and salt-and-peppers are depicted, representing the organization of neurons within V1. **c** The response shows neural activity at specific time frames (1ms to 100ms) after presenting the visual stimulus. Brighter spots indicate higher neural activity. The reconstruction shows the resulting images reconstructed from the neural responses at corresponding time frames. **d** The firing rates for both V1’s structures, and correspoding reconstructions are shown, with the pinwheel structures potentially exhibiting more firing wave patterns, while the salt-and-peppers show more randomness firing patterns.

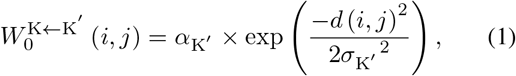

where K = E-or I-neurons; *i* = 1, 2, …, *N* (E- and I-neuron ID), the meaning and the value of the parameters can be found in Table 1, the same as below. Only the E-neuron receives stimuli from a corresponding area on the input image, based on its spatial location. To enable a fair comparison and consistent analysis, the SESNN module configured for forming different V1 features receives approximately the same total input image size. Moreover, the size and overlap of RFs are meticulously calculated to mirror biological consistency. Detailed parameter settings for these configurations can be found in Table 2.

**Table 1.**
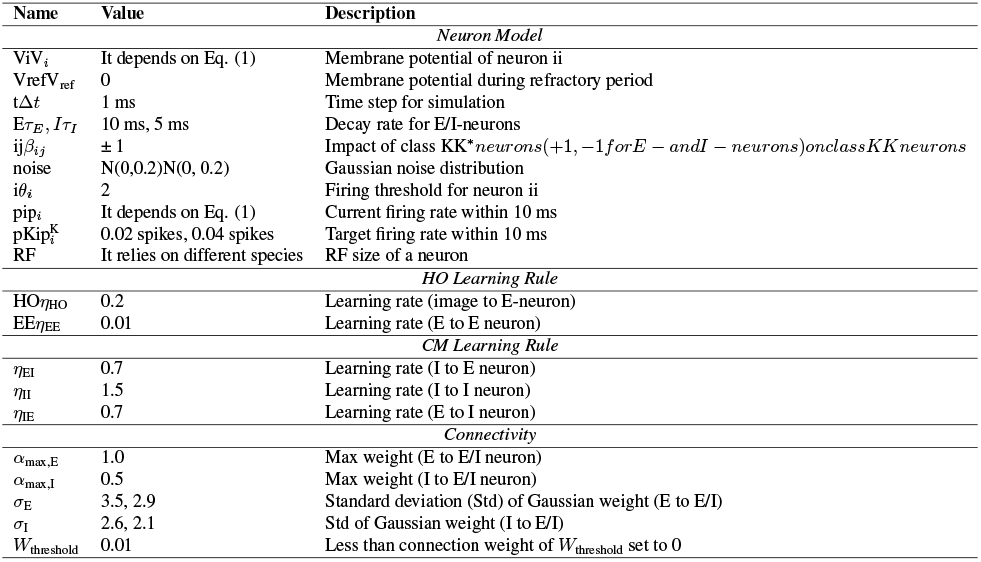
Neuron Model and Learning Rules Parameters.

**Table 2.**
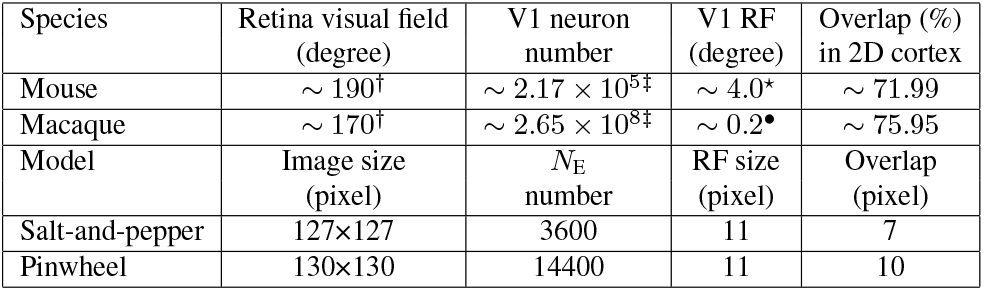
Comparative visual system metrics across species. *†* refer to [21] and [8]. *‡* refer to [26]. ⋆ refer to [22]. *•* [27].

The spiking behavior of neurons in the SESNN module is governed by the leaky integrate-and-fire (LIF) neuronal model, characterized by refractory periods of 3 ms and adaptive firing thresholds. The mathematical description of these dynamics is as follows:

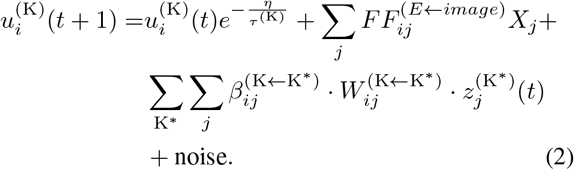

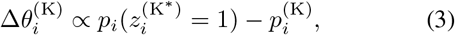

The membrane potential of each neuron decays over time, influenced by a time constant (10ms for E-neurons, 5ms for I-neurons), feedforward input from the natural image stimuli, recurrent input, and noise. When this potential exceeds a specific threshold, a spike occurs. Based on the discrepancy between the current and targeted firing rates, the firing threshold dynamically adapts.

In SESNN module, the synaptic weights are learned by two principal learning paradigms: Oja’s variant of the Hebbian Rule (HO) and the Correlation Measurement Rule (CM). These paradigms are instrumental after every 100 ms of the presentation of stimuli, changing the synaptic efficacies according to the following formula:

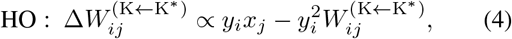

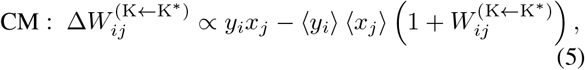

where *x, y* refer to the spike rate of presynaptic and postsynaptic neurons, the operator ⟨·⟩ denotes the lifetime average value.

Specifically, the HO rule reinforces synaptic connections upon the synchronicity of neuronal discharge and the CM rule preferentially augments the synaptic weights for neurons exhibiting correlated activity and attenuates those for neurons with anti-correlated activity. The model encapsulates distinct synaptic arrangements, wherein feedforward and recurrent connections between E-neurons are updated in accordance with the HO rule. Meanwhile, the CM rule is applied to govern the adjustment of other synaptic connections within the network.

The SESNN module generates a time-based raster plot of these rates, which we then use in conjunction with equation <u>6</u>for image reconstruction. The raster plot is transformed into a vector *r*, representing firing rates, where each neuron is associated with an RF. The reconstructed image, Img, is derived as a normalized weighted sum of these RFs:

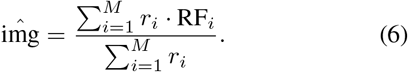

Here, *M* is the total number of neurons, *r*_*i*_ is the firing rate of the *i*-th neuron, and RF_*i*_ is the receptive field of the *i*-th neuron.

### 2.3 The DRUnet module

In the cerebral cortex, neuronal information processing predominantly occurs through generating action potentials, or ‘spikes.’ However, the mechanisms by which neurons in V1 encode complex visual stimuli through these spike patterns are not fully understood. Moreover, the distinctive characteristics of functional structures within V1 and their roles in encoding visual stimuli have not been comprehensively explored, particularly from a spike-based modeling perspective. This gap in knowledge necessitates a thorough investigation to uncover the complexities of neuronal encoding in V1.

While the SESNN module successfully reproduces various functional structures of V1 and allows for the recording of neuronal responses to natural image stimuli within a complete V1 framework, the precise mechanisms by which V1 neurons encode visual stimuli into individual spikes, and the differential functional roles of orientation structures in this process, remain elusive. To address these questions, we have developed a deep recurrent neural network based on the U-Net architecture (refer to Figure <u>2</u>f-h).

This network features a decoder composed of four segments: a transposed convolutional layer that converts neuronal population responses to an image-sized output, mirroring the inverse process of V1 neuronal stimulus reception. The kernel size of the transposed convolutional operation matches the neuron’s RF. Subsequently, the image enters a modified U-Net structure. Drawing inspiration from full convolutional networks commonly used in image reconstruction or segmentation, the first half of the U-Net involves convolutions and downsampling to extract signal features thoroughly. In contrast, the third layer employs a Convolutional Long Short-Term Memory (convLSTM) layer, designed to extract spatiotemporal information and the most abstract semantic content, believed to be related to past neuronal spike activity and the neuron’s encoding process based on prior knowledge.

In the second half of the network, transposed convolution upsamples the data, allowing for a broader representational space, followed by convolution and the recovery of a reconstructed representation of the image. Additionally, a recurrent neural network structure is employed to enable direct prediction of the information encoded by neurons at each time step, bypassing the need to recall information encoded in past spikes.

Two distinct output heads are implemented: one dedicated to the reconstruction of visual stimuli, representing the encoded images by V1 neurons, and the other focused on edge reconstruction. This latter aspect is crucial as it simulates a significant functional role of the pinwheel structure in edge detection. The architecture’s autoencoder segment incorporates skip connections, akin to those observed in U-Net. These connections facilitate the integration of both low-level features, which correspond to precise spatial location information in neuronal spikes, and high-level features, which are more aligned with the semantic content of the image, in the reconstruction process.

The learning objective function of the DRU-net is divided into four primary components, with the first being the Mean Squared Error (MSE) Loss. The MSE Loss is employed to train the reconstructed image head, aiming at the faithful reconstruction of the original image. This method is widely recognized and utilized in the field of computer vision.

To emulate the human visual system’s (HVS) ability to discern edges and highlight high-contrast details effectively, the pinwheel structure’s functionality is integrated into our model. This is achieved through the incorporation of the Structural Similarity Index (SSIM) into our loss function. SSIM, as formulated, evaluates the similarity between two images *x* and *y* based on their luminance, contrast, and structure, and is calculated as follows:

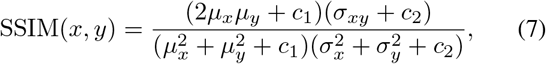

where *µ*_*x*_ and *µ*_*y*_ are the mean intensities, *σ*_*xy*_ is the covariance, and 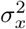 and 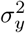 are the variances of *x* and *y* respectively, within the Gaussian window. We have employed a Gaussian filter with a window size of 11 to smooth each patch, and the SSIM thus computed is essentially a Mean-SSIM, averaged over these windows. Therefore, *µ* and *σ* in this context refer to the mean and variance calculated within each Gaussian window. The SSIM loss function, ℒ_*SSIM*_, is then defined as the cumulative sum of SSIM values calculated over all windows:

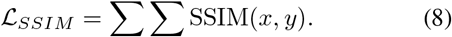

The Canny edge detector is utilized for edge detection from whitened images to guide the training of the reconstructed edge head of our model. The primary function of this edge head is to ascertain the likelihood of each pixel being an edge. To facilitate this, we employ a cross-entropy loss function, a standard approach in neural network-based classification tasks. Recognizing the disproportionate ratio of edge to non-edge pixels in typical images, we introduce a weighting factor in our loss function. This factor is designed to increase the significance of edge pixels, thereby addressing the imbalance and enhancing edge detection accuracy. The implementation process involves normalizing and applying Gaussian blurring to the original image. Subsequently, the Canny edge detector identifies the edges, which are then used as the teaching signal to train the neural network. The loss function is calculated as a class-balanced binary cross-entropy loss. The weighting factor is determined based on the average density of edge pixels in the detected edges. The loss function is expressed as follows:

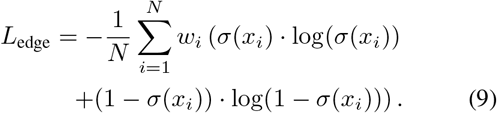

In addition to the primary loss components, our network incorporates an orthogonal regularization to maintain generality. This regularization enforces orthogonality in the weights used during convolution operations, a constraint we refer to as orthogonal loss ℒ_*orth*_. The total loss function for our network is thus composed of four distinct components: the primary loss ℒ_1_, the edge detection loss ℒ_*edge*_, the Structural Similarity Index (SSIM) loss ℒ_*SSIM*_, and the orthogonal loss ℒ_*orth*_. Mathematically, this can be expressed as:

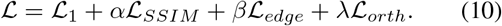

In our implementation, the coefficients *α*,*β*and *λ* are set to 1,1 and 0.0001, respectively, balancing the contribution of each loss component to the overall optimization process.

Previous studies have shown the viability of using a firing rate coding scheme for decoding information encoded by the primary visual cortex (V1) via deep neural networks[13, 28]. However, efforts to decode from a single time step perspective have been limited, highlighting the complexity of processing long time sequences in this context. In our approach, we leverage the firing rate decoding scheme as foundational knowledge. The training process involves two stages. Initially, we utilize DRUnet to train a decoder that interprets neuron firing rates over 100ms intervals. This training is conducted for 200 epochs, focusing on optimizing the network based on neuron firing patterns. Following this, we select the weight parameters that yield the best performance on the validation dataset, as determined by the loss function. These parameters are then employed to train a decoder that operates over 10 time steps, with each step incorporating 10ms of neuron firing rate data. Subsequently, the same methodology is applied to adapt DRUnet for decoding images directly from single time step spike data.

## 3 Results

### Firing patterns within pinwheels and salt-and-peppers

Our study, illustrated in Figure 2, uncovers a novel phenomenon in pinwheel structures named ‘firing waves.’ We observe that when a pinwheel processes a high-complex feature (see Figure 3), populational neurons, typically in PWCs, activate first. This initial response is sequentially followed by adjacent neurons, eventually spreading towards IODs. This distinctive firing pattern, undetected in previous experiments, is identified for the first time report in our SESNN module. Notably, such firing waves are absent in salt-and-pepper organizations. In the pinwheel structure, the reconstructed images gradually become more defined over time, suggesting a detailed and progressive encoding of the visual stimulus. For the salt-and-pepper structure, the reconstructions are less detailed, even at longer time frames, which aligns with the notion that this organization encodes the stimulus more rapidly but with less detail (refer to Figure 3).

**Figure 3.**
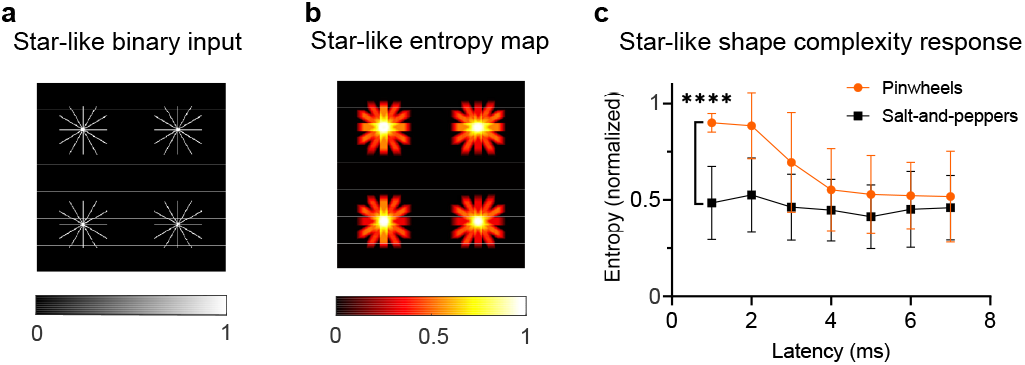
Comparison of the average complexity of first response time between pinwheels and salt-and-peppers. **a** Binary image containing multiple star-like shapes against a black background. **b** Entropy map corresponding to the binary input. **c** Comparing the response of two different neural populations: pinwheels and salt-and-peppers, over time (ms). All data: mean ± SD, \*\*\*\**p <* 0.0001.

### Trade-off between response speed and visual acuity in varying V1 functional structures

In nature, different species exhibit diverse survival requirements. For instance, mice necessitate rapid response times for evasion, prioritizing quick reactions over detailed visual processing. Conversely, higher animals like macaques benefit from precise visual acuity, aiding in complex environmental interactions. This variation suggests an evolutionary pressure leading to distinct V1 functional structures in different species. However, empirical evidence supporting this hypothesis has been limited.

Figure 3 illustrates the correlation between complexity and latency times in response to binary input (excluding the image contrast influence) for two cluster structures: pin-wheel structures and salt-and-pepper organizations. This figure highlights the sequential encoding process of image details by pinwheel—initially encoding complex details followed by other aspects. During this phase, the encoding of other details concurrently intensifies the orientations within complex details, thereby enhancing image acuity (sharp-ness) (refer to the fourth panel of <u>4</u>A.)

This comparison highlights a fundamental trade-off in V1 functional structures. The salt-and-pepper organization, akin to that observed in mice, offers rapid encoding at the expense of precision. On the other hand, the pinwheel structures, despite their longer encoding durations, excel in precision, attributable to their organized and coherent arrangement. This structured configuration in pinwheels facilitates a systematic and comprehensive response to visual stimuli, underscoring the adaptive evolutionary strategies in different species.

### Functional implications of pinwheel structures within V1 for edge detection

Our research delves into the critical characteristics that underlie the emergence of pinwheel structures in the visual cortex: the organization of receptive fields (RFs) and the influence of short-range lateral connections. We develop two targeted experiments to probe these features:

- **Pinwheel with shuffled RFs**: By randomizing the orientation preferences within the pinwheel structure, this model yields crucial insights into the impact of orientation preference clustering on functional outcomes.
- **Pinwheel with severed lateral connections**: In this configuration, the disruption of lateral connections between neurons eliminates the emergence of firing waves. This aspect offers a unique perspective on the role of interneuronal interactions in the propagation of firing waves.

Our comprehensive analysis employs a multidimensional approach. Utilizing the the proposed DRUnet model, we decode the neuronal spike activities across different firing patterns. The encoding quality of neurons is quantitatively assessed using metrics such as Mean Squared Error (MSE), Peak Signal-to-Noise Ratio (PSNR), Structural Similarity Index Measure (SSIM), and Local Phase Coherence (LPC)[7]. This study investigates four distinct pinwheel configurations to elucidate the functional significance of pinwheel connections and firing wave patterns.

Figure 4 examines the effects of local neuronal connectivity and RF arrangements on visual processing. The randomization of local connections diminishes detail clarity, indicating that such connections are integral for the rapid integration of visual information, especially within IODs, where neurons share similar orientation preferences. Additionally, the disruption of RF arrangements, which disrupts the IOD, leads to compromised edge continuity, signifying the IOD’s pivotal role in enhancing edge detail encoding.

**Figure 4.**
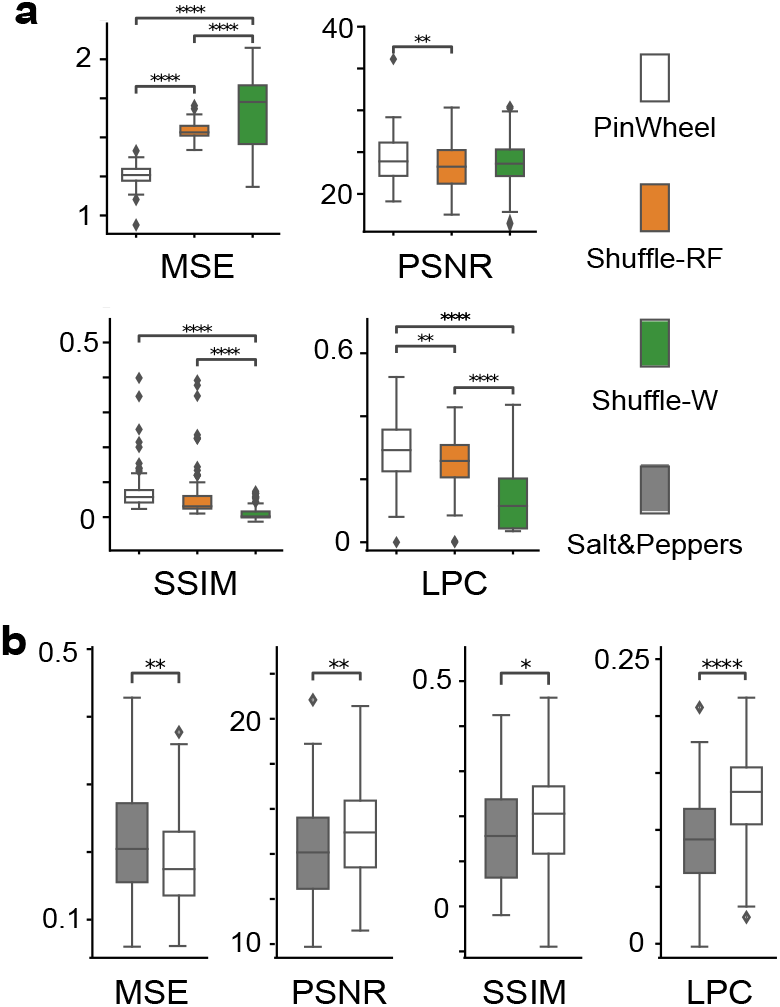
Different metrics for reconstruction quality assessment across pinwheels and salt-and-peppers ( * *p <* 0.05, ** *p <* 0.01, *** *p <* 0.001, **** *p <* 0.0001). **a** The results suggest that pinwheels generally have lower MSE, higher PSNR, and SSIM compared to the shuffled RF and connectivity within pinwheels. **b** Pinwheels improve more reconstruction accuracy and sharpness than salt-and-peppers.

The integrity of local lateral connections within pinwheels is paramount for encoding precision. The observed diffusion-like firing wave patterns of lateral connections facilitate the collective neural response to edge stimuli, underscoring the significance of synchronized activation patterns in edge encoding.

Conversely, when RFs of pinwheels are shuffled, encoding efficacy notably declines. This disruption disperses the orientation clusters, leading to a loss of functional coherence and, consequently, a less precise encoding. These findings highlight the necessity of structured orientation preferences for optimal neuronal encoding processes.

## 4 Discussion

This study has provided significant insights into the functional dynamics of V1, particularly emphasizing the roles of pinwheels and salt-and-peppers. Our proposed SESNN-DRUnet model, has led to several key discoveries:

1. The SESNN module has demonstrated remarkable efficacy in reconstructing pinwheel structures in a manner closely resembling biological reality. This capability allows for an in-depth exploration of the functional dynamics within V1, particularly in the context of orientation selectivity and spatial-temporal coding. The success of the SESNN module in accurately modeling these structures marks a significant advancement in our understanding of functional roles within pinwheel structures.

2. Coupled with the SESNN module, the DRUnet module has emerged as a potent tool for decoding the complex firing patterns within V1. This synergy has enabled us to delve deeper into the functional significance of firing waves, particularly in their role in enhancing orientation-specific processing. Our findings indicate that these firing waves, originating in pinwheel centers and propagating to IODs, contribute substantially to the detailed encoding of visual information, a critical aspect of sensory processing.

3. A critical trade-off between response speed and visual acuity in different V1 functional structures has been identified. Pinwheel structures, with their densely packed neurons and organized connectivity, exhibit a delayed yet detailed and precise encoding capability. This contrasts with the salt-and-pepper organizations, where a faster response time is achieved at the expense of reduced visual acuity. This discovery highlights the evolutionary adaptations in neural structures, balancing the need for rapid sensory processing against the demand for detailed and accurate visual representation.

Future studies should explore the link between initial responses in PCs and the creation of saliency maps within OPMs. This investigation could significantly enhance our comprehension of bottom-up processing in the visual system. Understanding how PCs’ early responses influence saliency map generation could reveal key mechanisms underlying the detection and prioritization of visually prominent features in the environment.

Our research thus presents a comprehensive understanding of the functional dynamics within V1, offering novel insights into the neural mechanisms underlying visual processing. The implications of these findings extend beyond basic neuroscience, providing valuable perspectives for the development of advanced image recognition algorithms in artificial intelligence and computational models that mimic biological visual processing.

## Notes

### Competing Interest Statement

The authors have declared no competing interest.

## References

[1] Amir Aghabiglou and Ender M Eksioglu. Projection-based cascaded u-net model for mr image reconstruction. Computer Methods and Programs in Biomedicine, 207:106151, 2021. 1

[2] Arish Alreja, Ilya Nemenman, and Christopher J. Rozell. Constrained brain volume in an efficient coding model explains the fraction of excitatory and inhibitory neurons in sensory cortices. 18(1):e1009642. Publisher: Public Library of Science. 2

[3] Anthony J. Bell and Terrence J. Sejnowski. The “independent components” of natural scenes are edge filters. Vision Research, 37(23):3327–3338, 1997. 1

[4] Yi Cai and Jiangying Yuan.A review of u-net network medical image segmentation applications. In Proceedings of the 2022 5th International Conference on Artificial Intelligence and Pattern Recognition, pages 457–461, 2022. 1

[5] Afshin Dehghan, Syed Zain Masood, Guang Shu, Enrique Ortiz, et al. View independent vehicle make, model and color recognition using convolutional neural network. arXiv preprint arXiv:1702.01721, 2017. 2, 1

[6] Ning Han, Li Zhou, Zhengmao Xie, Jingli Zheng, and Liuxin Zhang. Multi-level u-net network for image super-resolution reconstruction. Displays, 73:102192, 2022. 1

[7] Rania Hassen, Zhou Wang, and Magdy M. A. Salama. Image sharpness assessment based on local phase coherence. IEEE Transactions on Image Processing, 22(7):2798–2810, 2013. 7

[8] Christopher P. Heesy. On the relationship between orbit orientation and binocular visual field overlap in mammals. 281A(1):1104–1110. eprint: 10.1002/ar.a.20116. 3

[9] Stephen D. Van Hooser, J. Alexander F. Heimel, Sooyoung Chung, Sacha B. Nelson, and Louis J. Toth. Orientation selectivity without orientation maps in visual cortex of a highly visual mammal. 25(1):19–28. Publisher: Society for Neuroscience Section: Behavioral/Systems/Cognitive. 1

[10] D. H. Hubel. Single unit activity in striate cortex of unrestrained cats. The Journal of Physiology, 147(2):226–238, 1959. 1

[11] D. H. Hubel and T. N. Wiesel. Receptive fields, binocular interaction and functional architecture in the cat’s visual cortex. The Journal of Physiology, 160(1):106–154, 1962. 1

[12] Jaeson Jang, Min Song, and Se-Bum Paik. Retino-cortical mapping ratio predicts columnar and salt-and-pepper organization in mammalian visual cortex. 30(10):3270–3279.e3. Publisher: Elsevier. 1

[13] William F Kindel, Elijah D Christensen, and Joel Zylberberg. Using deep learning to probe the neural code for images in primary visual cortex. Journal of vision, 19(4):29–29, 2019. 5

[14] Teuvo Kohonen. Self-organized formation of topologically correct feature maps. 43(1):59–69. 1

[15] Margaret I. Law, Kathleen R. Zahs, and Michael P. Stryker. Organization of primary visual cortex (area 17) in the ferret. 278(2):157–180. eprint: 10.1002/cne.902780202.

[16] Ming Li, Xue Mei Song, Tao Xu, Dewen Hu, Anna Wang Roe, and Chao-Yi Li. Subdomains within orientation columns of primary visual cortex. 5(6):eaaw0807. Publisher: American Association for the Advancement of Science. 1

[17] Ziwei Liu, Ping Luo, Xiaogang Wang, and Xiaoou Tang. Deep learning face attributes in the wild. In Proceedings of International Conference on Computer Vision (ICCV), 2015. 2, 1

[18] Ilya Loshchilov and Frank Hutter. Decoupled weight decay regularization. arXiv preprint arXiv:1711.05101, 2017. 2

[19] S. Maji, J. Kannala, E. Rahtu, M. Blaschko, and A. Vedaldi. Fine-grained visual classification of aircraft. Technical report, 2013. 1

[20] Fan Min, Linrong Wang, Shulin Pan, and Guojie Song. D 2 unet: Dual decoder u-net for seismic image super-resolution reconstruction. IEEE Transactions on Geoscience and Remote Sensing, 61:1–13, 2023. 1

[21] Sohrab Najafian, Erin Koch, Kai Lun Teh, Jianzhong Jin, Hamed Rahimi-Nasrabadi, Qasim Zaidi, Jens Kremkow, and Jose-Manuel Alonso. A theory of cortical map formation in the visual brain. 13(1):2303. Number: 1 Publisher: Nature Publishing Group. 3

[22] Cristopher M. Niell and Michael P. Stryker. Highly selective receptive fields in mouse visual cortex. 28(30):7520–7536. Publisher: Society for Neuroscience Section: Articles. 3

[23] Bruno A. Olshausen and David J. Field. Sparse coding with an overcomplete basis set: A strategy employed by V1? Vision Research, 37(23):3311–3325, 1997. 1

[24] Xingjian Shi, Zhourong Chen, Hao Wang, Dit-Yan Yeung, Wai-Kin Wong, and Wang-chun Woo. Convolutional lstm network: A machine learning approach for precipitation nowcasting. Advances in neural information processing systems, 28, 2015. 1

[25] Xue Mei Song, Ming Li, Tao Xu, Dewen Hu, and Anna Wang Roe. Precise targeting of single microelectrodes to orientation pinwheel centers. 10(11):e3643. 1

[26] Shyam Srinivasan, C. Nikoosh Carlo, and Charles F. Stevens. Predicting visual acuity from the structure of visual cortex. 112(25):7815–7820. Publisher: Proceedings of the National Academy of Sciences. 3

[27] Edward J. Tehovnik and Warren M. Slocum. Phosphene induction by microstimulation of macaque v1. 53(2):337–343. 3

[28] Cem Uran, Alina Peter, Andreea Lazar, William Barnes, Johanna Klon-Lipok, Katharine A Shapcott, Rasmus Roese, Pascal Fries, Wolf Singer, and Martin Vinck. Predictive coding of natural images by v1 firing rates and rhythmic synchronization. Neuron, 110(7):1240–1257, 2022. 5

[29] William E. Vinje and Jack L. Gallant. Sparse Coding and Decorrelation in Primary Visual Cortex during Natural Vision. Science, New Series, 287(5456):1273–1276, 2000. 1

[30] Joel Zylberberg, Jason Timothy Murphy, and Michael Robert DeWeese. A sparse coding model with synaptically local plasticity and spiking neurons can account for the diverse shapes of v1 simple cell receptive fields. PLoS computational biology, 7(10):e1002250, 2011. 2

