## Supplemental material for "The Functional Role of Pinwheel Topology in the Primary Visual Cortex of High-Order Animals for Complex Natural Image Representation"

Supplementary Material

### 1. Abbreviation

- **V1**: Primary Visual Cortex
- **HO**: Hebbian Oja Rule
- **CM**: Correlation Measuring Rule
- **SESNN**: Self-evolving Spiking Neural Network
- **OPMs**: Orientation Preference Maps
- **IODs**: Iso-Orientation Domains
- **PCs**: Pinwheel Centers
- **DRUNet**: Deep Recurrent U-Net
- **CMF**: Cortical Magnification Factor
- **SSIM**: Structural Similarity Index
- **MSE**: Mean Squared Error
- **LPC**: Local Phase Coherence
- **RF**: receptive field

### 2. Glossary

- **Iso-orientation domain**: An iso-orientation domain refers to a region of neurons that prefer the same visual stimuli orientation, as depicted in Figure 1(c).
- **Pinwheel center**: Pinwheel centers exhibit a continuous change in preferred orientation around the singularity. The orientation preferences cyclically shift by 0-180 degrees in either a clockwise or counterclockwise direction along the singularity, as depicted in Figure 1(b) at point 1.
- **Saddle point**: In the vicinity of saddle points, the orientation preferences of neurons remain relatively equivalent, as depicted in Figure 1(b) at point 2. The neurons in this region tend to maintain similar orientation preferences.
- **Fracture**: A fracture refers to a linear region characterized by rapid changes in orientation preferences, as Figure 1(b) illustrates at point 3.
- **Linear zone**: The contours of the linear zone exhibit parallelism, indicating that the orientation changes linearly along a specific direction. This linear relationship is illustrated in Figure 1(b) at point 4, where the orientation preferences align along parallel lines.
- **Nearest neighbor distance**: The nearest neighbor distance refers to the distance between a pinwheel center and its closest neighboring pinwheel center.
- **Hypercolumn**: A hypercolumn refers to a region within the primary visual cortex that exhibits periodicity in the organization of the orientation map. It is characterized by a repeating pattern of orientation preferences, where neighboring neurons within the hypercolumn exhibit similar or related preferred orientations.

### 3. Technical Settings

#### 3.1. DRUnet Module

As briefly introduced in Section 2.1, the DRUnet is trained using a custom dataset. To enhance the generalizability of DRUnet in decoding various scenes and to leverage high-quality images for improved decoding accuracy, we employed images from three distinct datasets:

- Portraits from the CelebA dataset [17], utilizing the first 4000 images.
- Automobile images from the Stanford Car Dataset [5], where the training set is alphabetically ordered, and the first 4000 images are selected.
- Aircraft images from Maji et al. [19], also using the first 4000 images.

The aforementioned data were amalgamated to form a new dataset. All images were whitened using the same procedure as those presented to the SESNN module. The processed dataset was then exposed to the SESNN module for 100 ms, during which all neuronal spike trains were collected. Subsequently, the paired image and neuronal response data were randomly divided into training and testing sets for the DRUnet.

In training the DRUnet module, images were clipped to the range (-1,1) without normalization. A Canny edge detector was employed for edge detection, setting the low and high thresholds at 20 and 50, respectively.

#### 3.2. Implementation Details

We implemented two DRUnet modules to decode images from pinwheel and salt-and-pepper SESNN modules. The DRUnet is an adapted version of the U-net, a model extensively employed in image reconstruction tasks.[1, 4, 6, 20] These modules share a similar architecture, with modifications to accommodate varying neuron population sizes. The initial layer in both architectures is a ConvTrans layer. It receives neuron spikes (neuron firing rates) as inputs and emulates neuron receptive fields, converting images from neuron population dimensions to their original sizes. The kernel size and stride correspond to the receptive field size and its overlap, with the channel count set to 3. Specifically, for pinwheel and salt-and-pepper setups, the kernel size is 12, with overlaps of 3 and 5, respectively.

Following layers include two downsample blocks and a ConvLSTM downsample block. Each consists of three convolution operations with ReLU activation and batch normalization. The ConvLSTM block [24] features a

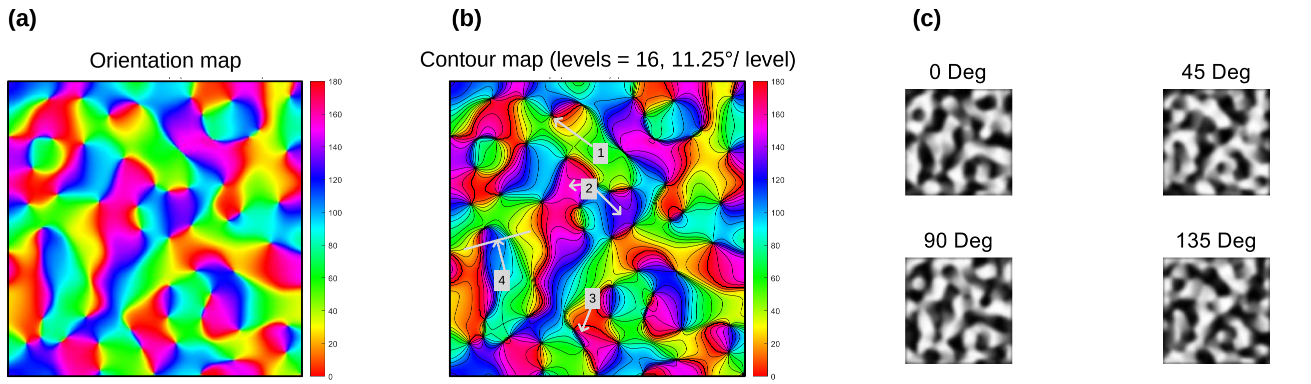


Figure 1. Orientation map, its contour map, and iso-orientation domains. (a) Orientation map generated by our SE-SNN model. (b) Point 1: pinwheel center. Point 2: saddle points. Point 3: fracture. Point 4: linear zone. (c) 0, 45, 90, and 135 degrees iso-orientation domains (white parts) within the orientation map.

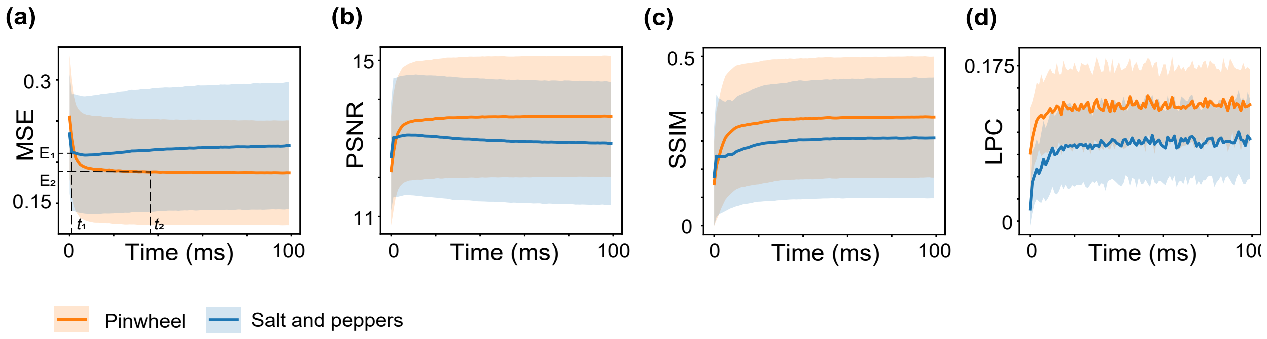


Figure 2. Comparative dynamics in reconstructions between pinwheel and salt-and-pepper organizations. **a** It shows the Mean Squared Error (MSE) of reconstructions for both pinwheel structures and salt-and-pepper organizations across time. **b** Peak Signal-to-Noise Ratio (PSNR) ascends over time for pinwheels and salt-and-peppers, implying progressively refined reconstructions. **c** Structural Similarity Index (SSIM), which measures image quality based on perceived changes in structural information, contrast, and luminance, also shows an increase over time. **d** Local Phase Coherence (LPC) scores, indicating the acuity of reconstruction between pinwheels and salt-and-peppers. Across panels **a-d**, solid lines denote the mean, while shaded areas indicate the standard deviation.

32-dimension hidden layer, replacing the initial convolution operation. Convolution operations have kernel sizes, strides, and paddings of (3,1,1), (3,1,1), and (3,2,1). The output from the second convolution is integrated with deeper features in a U-Net-like architecture.

The upsampling layer, reminiscent of the U-Net architecture, comprises three upsampling blocks, each available in two distinct versions. Each block consists of two convolution operations followed by a convolution transpose operation. The convolution operations maintain consistent dimensions with (3,1,1) and (3,1,1), respectively. The convolution transpose operation, however, varies with (3,2,1) and includes alternate out-paddings of 0 or 1. This configuration allows for precise image resizing, accommodating even odd-sized original images. A unique feature of this architecture is the recurrent connection in the DRUnet, where the output of the last upsampling layer is concatenated back to its input, enhancing the network's capability to preserve and integrate detailed image features across layers.

To decode the deep features outputted by the DRUnet, two convolution blocks are employed: one for image reconstruction and another for edge detection. Both blocks share a similar structure: three convolution layers with dimensions (3,1,1). Each layer employs a tanh activation function and batch normalization, except for the final layer of the edge detection block, which omits the tanh activation and the final layer of both omits batch normalization.

During the training phase, the DRUnet was optimized using the AdamW algorithm [18], starting with an initial learning rate of 0.01. The DRUnet was trained over 200 epochs to decode firing rates, using the best validation set performance to further train the network on 10-step time sequences (each step representing 10ms of firing rate) for an additional 200 epochs. The final training phase involved decoding single time steps for 200 epochs, again using the best validation performance. All training procedures were conducted on an Nvidia A100 Tensor Core GPU.

### 4. Trade-off between coding time and acuity within V1 organizations

Pinwheels show a steady approach to a stable MSE (Figure 2(a)), suggesting a lengthier coding time ($t_2 > t_1$) but more precise details ($E_2 < E_1$), whereas salt-and-peppers stabilize more rapidly yet with reduced precision. It is noteworthy that the curve representing the salt-and-pepper structures shows an uptick after the completion of the reconstruction, which can be attributed to noise disturbance. PSNR ascends over time for both types (Figure 2(b)), implying progressively refined reconstructions. SSIM increases (Figure 2(c)), reflecting heightened accuracy in visually representing the two V1 structures. LPC reveals that pinwheels achieve a higher acuity of reconstruction than salt-and-peppers (Figure 2(d)), which display lower LPC values.

The overall trend across these metrics of MSE, PSNR, SSIM, and LPC suggests that the model is more effective in reconstructing the pinwheel structures compared to the salt-and-peppers. The pinwheel structures demonstrate a delayed but more stable convergence in MSE (coding time) and achieve higher LPC scores (visual acuity), indicative of a more accurate and coherent reconstruction.

### 5. Code Availability

In accordance with the principles of open science and reproducibility, the complete codebase associated with this study will be made available in a public repository immediately upon the acceptance of this article for publication.
